# Mycorrhizas shape the evolution of plant adaptation to drought

**DOI:** 10.1101/2022.05.13.491064

**Authors:** Marco Cosme

## Abstract

- Plant adaptation to drought facilitates major ecological transitions, and is likely to play a vital role under looming climate change. Mycorrhizas can influence the physiological capacity of plants to tolerate drought. Here, I show how mycorrhizal strategy and drought adaptation shape one another throughout the course of plant evolution.
- To characterize the evolutions of both plant characters, I applied a phylogenetic comparative method using data of 1,638 extant species globally distributed.
- The detected correlated evolution unveiled gains and losses of drought tolerance occurring at faster rates in lineages with an ecto- or ericoid mycorrhizal strategy, which were on average about 15 and 300 times quicker than that in lineages with the arbuscular mycorrhizal and naked root (non-mycorrhiza or facultatively arbuscular mycorrhiza) strategy, respectively. Among mycorrhiza shifts, the arbuscular mycorrhiza losses in drought sensitive lineages were more frequent than any symbiont switching or other mutualism breakdown.
- My study suggests that mycorrhizas play a key facilitator role in the evolutionary process of plant adaptation to critical changes in water availability across global climates.

## Introduction

The adaptation of plants to contrasting gradients of water availability has facilitated major ecological transitions on Earth, such as the migration from exclusively aquatic environments to terrestrial habitats (Rubinstein *et al*., 2010; Oliver *et al*., 2020), and the subsequent expansions across nearly all land surfaces (Joswig *et al*., 2021), of which about half are currently susceptible to droughts (Mishra & Singh, 2010). These worldwide evolutionary adaptations in plants have coexisted for millions of years with strategic associations between plant roots and mycorrhizal fungi (Redecker *et al*., 2000; Brundrett & Tedersoo, 2018a; Oliver *et al*., 2020), which influence the drought tolerance and survival of extant plant species (Augé, 2001; Lehto & Zwiazek, 2011). However, how plant adaptation to drought and mycorrhizal strategy influence one another throughout evolution is unknown.

Droughts occur in nearly all climatic zones, from high to low rainfall areas, and result primarily from a reduction in precipitation below the normal levels over an extended period of time, such as a season or a year (Mishra & Singh, 2010). These events can lead to ecosystem instability and reordering, disturbed riparian habitats, reduced pasture productivity, and crop failure (Mishra & Singh, 2010; Ploughe *et al*., 2019; Püschel *et al*., 2021). Paleo records suggest that drought events led to significant vegetation shifts in a distant past (Schmieder *et al*., 2013; Clifford & Booth, 2015; Martínez-Vilalta & Lloret, 2016). In recent years, droughts have been experienced with higher peaks and severity due to climate change (Mishra & Singh, 2010), and already caused a few vegetation shifts regardless of climate or vegetation types (Martínez-Vilalta & Lloret, 2016). Current projections indicate that future global warming will lead to increased manifestations of droughts in regions where drought intensification does not yet occur (Ploughe *et al*., 2019; Zittis *et al*., 2019; Spinoni *et al*., 2020), which is likely to alter the regimes of natural selection.

Plants, as the main primary producers that fixate carbon and provide shelter and food for a myriad of organisms, are essential to most land ecosystems across climatic zones, and are on the frontline of perilous droughts. Drought disrupts water relations in plants when the rate of transpiration exceeds water uptake, causing drought stress, to which plants deploy adaptive responses that tend to be species-specific (Ploughe *et al*., 2019; Bowles *et al*., 2021). The responses that allow plants to buffer against periods of drought stress (e.g. stomatal closure, root growth) can be collectively defined as mechanisms of tolerance, while those that result in chronic physiological disruption or mortality are symptomatic of sensitivity (Ploughe *et al*., 2019; Bowles *et al*., 2021). Our current knowledge on how plant tolerance and sensitivity to drought evolve is extremely limited. To my knowledge, only one study has investigate this across vascular plants based on 178 extant species, of which more than two thirds have domesticated traits (Bowles *et al*., 2021). Hence, data from a large diversity of species living in nature across the globe is critically needed to narrow down uncertainties when characterizing the evolution of plant adaptation to drought in terrestrial ecosystems.

Paleo records, evolutionary history, and current distribution strongly indicate that the large majority of Earth’s natural vegetation has evolved in association with mycorrhizal fungi (Remy *et al*., 1994; Redecker *et al*., 2000; Brundrett & Tedersoo, 2018a; Soudzilovskaia *et al*., 2019). These associations are strategically important for plants because they provide multiple benefits at a relatively low cost in terms of spent carbon (Smith & Read, 2008). These benefits comprise the uptake, transport, and delivery of mineral nutrients by fungal mycelia across the soil into the roots, as well as the improvement of plant growth, survival, and protection. The most ubiquitous classes of mycorrhizas are the arbuscular, ecto-, ericoid, and orchid mycorrhiza, which are primarily distinguished based on host plant and characteristic symbiotic structures (Van der Heijden *et al*., 2015; Brundrett & Tedersoo, 2018a; Miyauchi *et al*., 2020). Plants that generally do not form mycorrhizas, and rely therefore more on their own roots, are non-mycorrhizal, while those that can form mycorrhizas in certain conditions but remain non-mycorrhizal in many others are facultatively mycorrhizal (Cosme *et al*., 2018; Lambers *et al*., 2018; Soudzilovskaia *et al*., 2020). Despite these primary distinctions, the ecto- and ericoid mycorrhizas use a similar strategy for taking up nutrients, occupy niches with relatively similar characteristics, and even share some of the same fungal symbionts (Peat & Fitter, 1993; Van der Heijden *et al*., 2015; Gerz *et al*., 2018; Lu & Hedin, 2019). Furthermore, the ecto-, ericoid, and arbuscular mycorrhizas have in common the ability to influence the physiological capacity of their host plants to tolerate drought, while drought can also influence the development of these mycorrhizas (Augé, 2001; Lehto & Zwiazek, 2011; Sebastiana *et al*., 2018; Kilpeläinen *et al*., 2020; Mu *et al*., 2021; Püschel *et al*., 2021). A major gap of knowledge is still how plant adaptation to drought and mycorrhizas influence one another at an evolutionary time scale.

Several important aspects of plant evolution has been previously investigated with great efficiency, rigor, and objectivity using the hidden Markov model (HMM) in its binary trait-based approach, including the study of symbiotic nitrogen fixation and mutualism breakdown (Beaulieu *et al*., 2013; Werner *et al*., 2014; Werner *et al*., 2018; Joly & Schoen, 2021). Here, I used the recently developed multistate HMM approach, which arguably increases the accuracy of estimations along deep phylogenies (Boyko & Beaulieu, 2021), to analyze the relationship between plant adaptation to drought and mycorrhizal strategy measured across a global sample of hierarchically evolved species. To this end, I collected species’ data from several large published datasets and ran a series of HMMs of incrementing complexity to test the hypotheses that: (1) the evolutions of these two functionally important plant characters depend on each other; and (2) mycorrhizas markedly influence the speed of evolution of drought adaptation in plants. Finally, I analyzed the sensitivity of main conclusions in relation to data uncertainty.

## Materials and methods

### Data on plant adaptation to drought

I downloaded trait data on ‘species tolerance to drought’ (TraidID 30) for 3,324 plant species from the Try – Plant Trait Database (Kattge *et al*., 2020), distributed among seven original datasets. This trait was renamed hereafter as ‘drought adaptation’ for a clear distinction between trait name and state. From the original datasets obtained, only the larger four (DatasetID 49, 68, 92, and 98) showed significant species’ overlaps with the FungalRoot dataset (the larger dataset used here to assign the mycorrhizal strategy, as described below). Because DatasetID 49 and 68 were remarkably identical, with the former being slightly larger and accompanied by detailed information on data determination, only the three original datasets DatasetID 49, 92, and 98 were used in this study (Niinemets & Valladares, 2006; Green, 2009; Wirth & Lichstein, 2009).

To standardize data interpretation across datasets, and facilitate model estimation of parameters, the drought adaptation was simplified into a binary variable, with species having either a ‘tolerant’ or ‘sensitive’ state, following a previous approach (Bowles *et al*., 2021). For DatasetID 49, the original values varied continuously between 0 and 5, with 1, 2, 3, 4, and 5 standing for ‘very intolerant’, ‘intolerant’, ‘moderately tolerant’, ‘tolerant’, and ‘very tolerant’ to drought, respectively (Niinemets & Valladares, 2006). Therefore, species with original value of less than 2.5 were assigned here with a ‘sensitive’ state, whereas the remaining species were assigned with a ‘tolerant’ state. For DatasetID 92, the original values varied categorically among ‘none’, ‘low’, ‘medium’, and ‘high’ tolerance to drought. Hence, the species with the original value ‘none’ or ‘low’ were assigned here with a ‘sensitive’ state, whereas the species with original value ‘medium’ or ‘high’ were assigned with a ‘tolerant’ state. For DatasetID 98, the original values varied categorically among ‘none, dies off in dry conditions’, ‘medium, dies off after several months’, ‘fairly drought resistant’, and ‘very drought resistant’. Thus, the species with the original value ‘none, dies off in dry conditions’ or ‘medium, dies off after several months’ were assigned here with a ‘sensitive’ state, whereas the species with original value ‘fairly drought resistant’ or ‘very drought resistant’ were assigned with a ‘tolerant’ state.

Although this approach allowed state standardization across datasets, it still led to assignment mismatches for some species, generating data uncertainty (addressed below in sensitivity analysis). This occurred only for the assignments using DatasetID 92 against that of 49 or of 98. However, by considering these mismatches, I assembled dataset versions with partial differences of plant adaptation to drought by prioritizing the assignments based either on DatasetID 92 or on both DatasetID 49 and 98 together (described below in assembly of dataset versions).

### Data on plant mycorrhizal strategy

Data on plant mycorrhizal types were obtain from two sources, the FungalRoot database (Soudzilovskaia *et al*., 2020) and an alternative dataset (Bueno *et al*., 2018; hereafter Bueno dataset). The FungalRoot is the largest database of its kind ever assembled, containing mycorrhizal assignments for 14,347 plant genera. The main mycorrhizal types included in the FungalRoot are the arbuscular mycorrhizal (AM), facultatively AM, ecto-mycorrhizal, dual ecto-mycorrhizal and AM, ericoid mycorrhizal, orchid mycorrhizal, and non-mycorrhizal, which follow the mycorrhiza definitions previously proposed (Brundrett, 2009; Brundrett & Tedersoo, 2018a). However, because these definitions are still in part a matter of debate (Brundrett & Tedersoo, 2018b; Bueno *et al*., 2018; Bueno *et al*., 2019; Brundrett, 2021; Bueno *et al*., 2021), to address the issue of data uncertainty, I generated partially different dataset versions on mycorrhizal strategy using the relatively smaller Bueno dataset (described below in assembly of dataset versions). The Bueno dataset includes assignments of AM, ecto-mycorrhiza, ericoid mycorrhiza, and non-mycorrhiza for a total of 1,362 plant species.

Similar to the data on drought adaptation, to facilitate model estimation of parameters, the mycorrhizal strategy was simplified here into a ternary variable following a previous approach (Lu & Hedin, 2019). Central to this approach is the observation that plants have evolved different strategies for investing photosynthetically fixed carbon to compete for limiting soil resources: (1) by scavenging for plant-available nutrients mainly in symbiosis with AM fungi (Smith & Smith, 2011; Lu & Hedin, 2019) (hereafter ‘AM strategy’); (2) by mining organic-bound nutrients primarily in symbiosis with ecto- or ericoid mycorrhizal fungi (Talbot *et al*., 2008; McGuire *et al*., 2010; Lu & Hedin, 2019) (hereafter ‘EEM strategy’); or (3) by taking up resources mostly via the absorptive surface of their own naked roots (Brundrett & Tedersoo, 2018a; Cosme *et al*., 2018; Lu & Hedin, 2019) (hereafter ‘NR strategy’). In addition, monophyletic mycorrhizas were not considered here alone to minimize phylogenetic comparative issues related to single evolutionary transitions (Gardner & Organ, 2021). Thus, to determine the mycorrhizal strategy using the FungalRoot database, plant species belonging to genera with AM type were assigned here with an AM strategy, while those belonging to genera with either ecto- or ericoid mycorrhizal type were assigned with an EEM strategy. In addition, because many dual mycorrhizal plants tend to be dominated by AM during their seedling stage only (Brundrett & Tedersoo, 2018a; Teste *et al*., 2020), the plant species belonging to genera with dual ecto-mycorrhizal and AM type were considered here have a predominant EEM strategy. Furthermore, because many non-mycorrhizal and facultatively AM plants are habitat or nutritional specialists where mycorrhizas are described to be less relevant (Brundrett & Tedersoo, 2018a; Cosme *et al*., 2018), the plant species belonging to genera with either non-mycorrhizal or facultatively AM type were assigned here with a NR strategy. The only three detected plant species that belong to genera with the orchid mycorrhizal type were excluded from the analysis. To determine the mycorrhizal strategy using the Bueno dataset, the plant species with AM, ecto- or ericoid mycorrhizal, and non-mycorrhizal type were assigned here with the AM, EEM, and NR strategy, respectively.

### Plant phylogeny data

I obtained two plant phylogeny versions (GBMB and GBOTB) from Smith and Brown (2018), which is the most broadly inclusive, time-calibrated plant phylogeny construction published to date. Both phylogeny versions were constructed with GenBank taxa. However, GBMB has 79,874 taxa and backbone provided by Magallón *et al*. (2015), while GBOTB has 79,881 taxa and a backbone provided by the Open Tree of Life version 9.1 (Smith & Brown, 2018). These two phylogeny versions were analyzed here to account for potential phylogenetic uncertainty. To assign the phylogenetic relationships among species, both phylogenetic trees were pruned by keeping only the tips whose labels matched the names of the species included in the assembled datasets (described below in assembly of dataset versions).

### Assembly of dataset versions

Based on the data assignments described above, a total of six partially different dataset versions were assembled, hereafter named dataset v1 to v6: *dataset v1* consists of the adaptation to drought prioritizing DatasetID 92 of TRY, mycorrhizal strategy based on FungalRoot database, and phylogeny mapped against the GBMB tree; *dataset v2* consists of the adaptation to drought prioritizing DatasetID 92 of TRY, mycorrhizal strategy based on FungalRoot database, and phylogeny mapped against the GBOTB tree; *dataset v3* consists of the adaptation to drought prioritizing DatasetID 49 and 98 of TRY, mycorrhizal strategy based on FungalRoot database, and phylogeny mapped against the GBMB tree; *dataset v4* consists of the adaptation to drought prioritizing DatasetID 49 and 98 of TRY, mycorrhizal strategy based on FungalRoot database, and phylogeny mapped against the GBOTB tree; *dataset v5* consists of the adaptation to drought prioritizing DatasetID 92 of TRY, mycorrhizal strategy based on Bueno dataset, and phylogeny mapped against the GBMB tree; and *dataset v6* consists of the adaptation to drought prioritizing DatasetID 92 of TRY, mycorrhizal strategy based on Bueno dataset, and phylogeny mapped against the GBOTB tree. The dataset v1 to v4 had 1,638 species each, while dataset v5 and v6 had 787 species each. A summary of the dataset assembly is provided in Supporting Information Table S1 and the files of the respective phylogenetic trees and data frames are provided in Supporting Information Data S1 (found at https://github.com/marcosmeweb/symbio-inc-drought). These dataset versions were analyzed separately and then used to generate average results and to characterize the impact of data uncertainty on main conclusions (described below in sensitivity analysis).

### Global mapping of species distribution

I downloaded the georeferenced occurrences (latitude and longitude) for vascular plant species from the Global Biodiversity Information Facility (GBIF.org, 23 December 2021), which included 457,547 occurrences distributed among 25,779 species. Then, to validate the global scale of my analysis, I used this georeferenced data to assign occurrences to the maximum number of species included in the assembled dataset versions (Supporting Information Data S2 found at https://github.com/marcosmeweb/symbio-inc-drought).

### Modeling of plant character evolution

To test for correlated character evolution and estimate evolutionary transition rates between states, I ran 18 hidden Markov models (HMMs) of incrementing complexity, using the corHMM function of the R package corHMM v2.1 (Boyko & Beaulieu, 2021). Briefly, the corHMM function takes a phylogenetic tree and state data to estimate among several parameters the transition rates among states of discrete characters. The models tested here included either a dependent or independent mode of evolution for the characters ‘drought adaptation’ and ‘mycorrhizal strategy’. Moreover, the corHMM function allows to detect the occurrence of unobserved (or hidden) phylogenetic factors that have either promoted or constrained the evolutionary processes of the observed characters, controlling for phylogenetic bias (Beaulieu *et al*., 2013; Boyko & Beaulieu, 2021). Thus, each model tested here included either one, two, or three categories of hidden rates. Moreover, each model included a structure of either homogeneous evolution (all evolutionary transition rates are equal; ER), partially homogenous evolution (only evolutionary transition rates between any two character states do not differ; SYM), or heterogenous evolution (all evolutionary transition rates differ; ARD) (Boyko & Beaulieu, 2021). Each model was run with five replicated starts, resulting in a total of 810 independent optimization exercises (output files provided in Supporting Information Data S3 are found at https://github.com/marcosmeweb/symbio-inc-drought). The corHMM function was run in R v3.6.2 using a supercomputer cluster. The codes use to analyze each dataset version are provided in the Supporting Information Note S1 at https://github.com/marcosmeweb/symbio-inc-drought. Finally, I compared the sample size-corrected Akaike information criteria (AICc) among models to select the best fitted one, and obtained the respective estimated evolutionary transition rates among the plant character states.

### Sensitivity analysis

I analyzed the robustness of model fitness and of estimated evolutionary transition rates in relation to three primary sources of uncertainty: 1) adaptation to drought data; 2) mycorrhizal strategy data; and 3) phylogenetic tree backbone. To this end, each of the 18 HMMs (described above in modeling of plant character evolution) were run on each of the six dataset versions (Supporting Information Table S1), using five replicated starts each. The robustness of model selection was evaluated by comparing how the different dataset versions changed the model ranking based on the AICc. The robustness of the evolutionary transition rates was performed by visual comparison of how the different dataset versions changed the transition rate parameters estimated by the respective best fitted models.

### Determining the phylogenetic imbalance ratio

To avoid erroneous detection of correlated evolution due to limited evolutionary sample size and/or state phylogenetic imbalance, it is recommended that each dataset holds a phylogenetic imbalance ratio (PIR) below 0.1 (Gardner & Organ, 2021). Hence, I have calculated the PIR for each of the six dataset versions analyzed here using the following formula proposed by Gardner & Organ (2021):

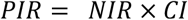

In this formula, *CI* is the consistency index and was calculated using the CI function in the R package “phangorn”, while *NIR* is the normalized imbalance ratio calculated using the following formula:

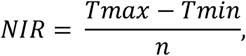

where *n* is the size of the dataset and *Tmax* and *Tmin* is the maximum and minimum frequency of a state, respectively.

## Results

### The evolutions of plant adaptation to drought and mycorrhizal strategy depend on each other

My first aim was to analyze the relationship between the two functionally important plant characters drought adaptation and mycorrhizal strategy measured across a global sample of hierarchically evolved species. To this end, I collected data from published trait, phylogeny, and occurrence databases of land plants (angiosperms and gymnosperms). Of the total number of plant species included in my analysis, 22%, 9%, and 8% were always considered to be drought sensitive and to have AM (arbuscular mycorrhizal), EEM (ecto- or ericoid mycorrhizal), and NR (naked root; i.e. non-mycorrhizal or facultatively AM) strategy, while 38%, 10%, and 3% were always considered to be drought tolerant and to have AM, EEM, and NR strategy, respectively (Fig. 1). The remaining 9% of the species had uncertain drought adaptation and mycorrhizal strategy depending on the assembled dataset version (Fig. 1; Supporting Information Table S2). Furthermore, the analyzed species belong to a total of 628 genera, 151 families, and 50 orders, representing a large diversity of terrestrial plants currently living in nature (Fig. 2a). To verify whether the species included in my analysis reflect a global sampling, I mapped a total of 112,276 geographical occurrences for 1,066 species (Supporting Information Data S2). The mapping of 65% of the species included in my analysis was sufficient to confirm that the majority is currently distributed across nearly all continents (Fig. 2b).

**Fig. 1.**
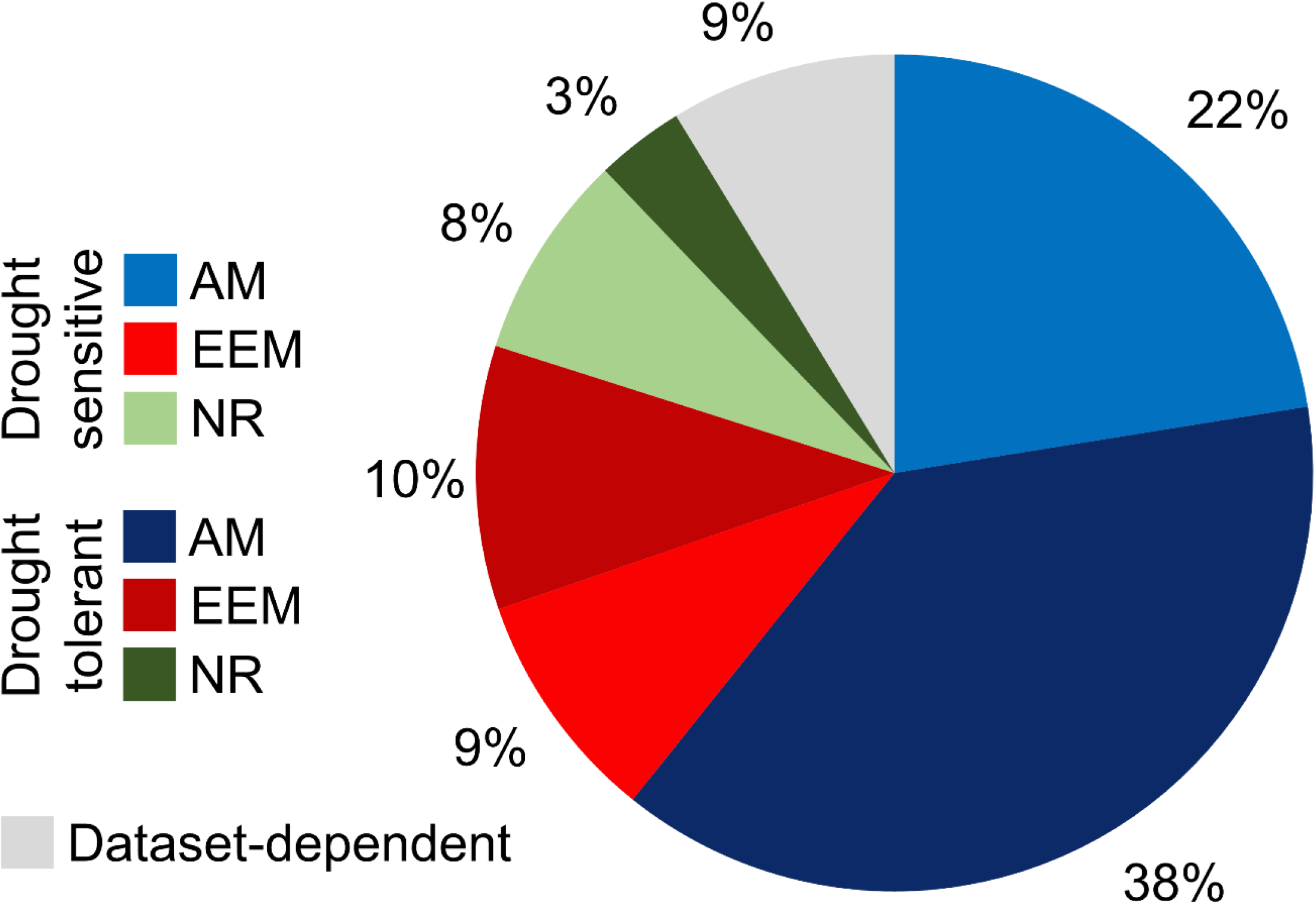
Percent distribution of mycorrhizal strategy and drought adaptation states. The pie chart includes 1,638 plant species (angiosperms and gymnosperms) colored according to their combined states of mycorrhizal strategy (AM, arbuscular mycorrhizal; EEM, ecto- or ericoid mycorrhizal; or NR, naked root, i.e. non-mycorrhizal or facultatively AM) and drought adaptation (sensitive or tolerant), or a dataset-dependent state. The dataset-dependent state corresponds to mycorrhizal strategy and drought adaptation assignments varying according to dataset v1 to v6 (for details on the assembly of dataset versions see materials and methods and Supporting Information Table S1).

**Fig 2.**
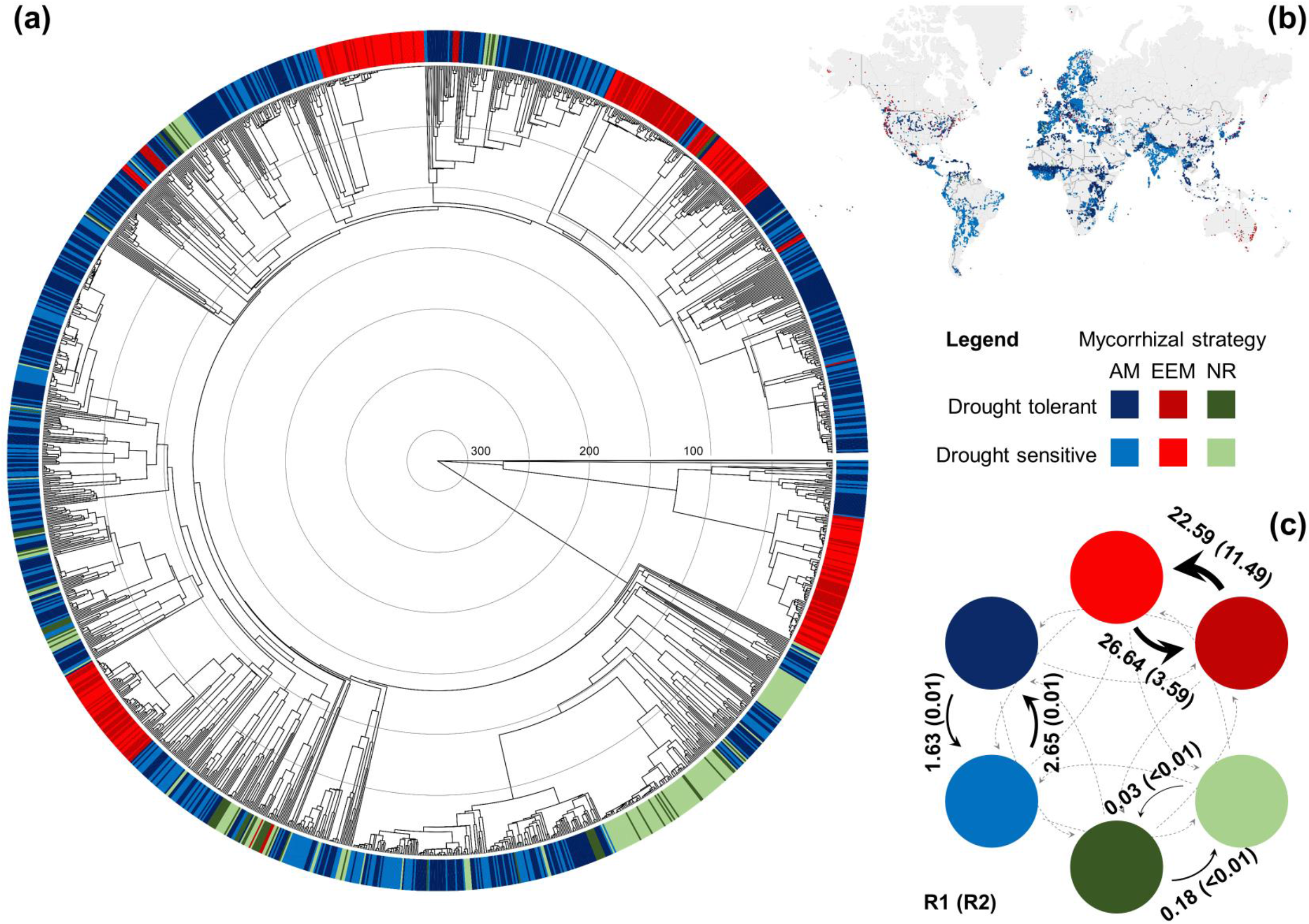
Phylogenetic and geographical distribution and transition rates of the dependent evolution of mycorrhizal strategy and drought adaptation. (a) Time-calibrated (Mya) phylogenetic tree of 1,638 plant species (angiosperms and gymnosperms) with a colored band at the tips indicating the mycorrhizal strategy (AM, arbuscular mycorrhizal; EEM, ecto- or ericoid mycorrhizal; or NR, naked root, i.e. non-mycorrhizal or facultatively AM) and drought adaptation (sensitive or tolerant) for each species as described in the legend and assigned according to dataset v1. The files of this and the other phylogenetic trees, with legible species labels, as well as the respective data frames, with mycorrhizal strategy and drought adaptation assignments, are provided in Supporting Information Data S1. The concentric circles inside the phylogenetic tree indicate 50 My. (b) Global geographical occurrences colored according to the mycorrhizal strategy and drought adaptation of 65% of the species included in the phylogenetic tree (for individual occurrences see Supporting Information Data S2). (c) Evolutionary transition rates (per 100 My) averaged from the estimates of the best fitted models of dependent evolution between mycorrhizal strategy and drought adaptation for dataset v1 to v6. For details on assembly of dataset versions see Supporting Information Table S1 and materials and methods. Values without and within brackets are transitions rates observed in the hidden rate category one (R1) and two (R2), respectively. Width of black arrows is proportional to the rates observed in R1. The dotted grey arrow indicates that the rate is < 0.01. The values of all the estimated evolutionary transition rates are provided in Table 2 and Supporting Information Table S3.

With the global sampling described above, I then tested whether the evolutions of plant adaptation to drought and mycorrhizal strategy are correlated with each other. This was done by comparing the fitness to the data among 18 HMMs with either an independent or dependent mode of evolution. These HMMs also differed among each other in term of hidden rate categories (from one to three) and evolution model structure (from fully homogenous to fully heterogenous). After a total of 810 optimization exercises, the resulting AICc of the HMMs provided strong statistical support for a dependent evolution between the plant characters drought adaptation and mycorrhizal strategy (Table 1), confirming my first hypothesis. This signifies that, throughout the course of plant evolution, the rate of change in drought adaptation (shifts between sensitive and tolerant) in a given lineage strongly depends on the mycorrhizal strategy (whether AM, EEM, or NR) of that lineage, and the rate of change in mycorrhizal strategy (shifts among AM, EEM, and NR) strongly depends on the lineage adaptation to drought (whether sensitive or tolerant) (Table 2). In addition, the best fitted HMMs revealed that this correlated evolution was influenced by an unobserved (hidden) phylogenetic factor with two state levels (Table 1), which has either promoted or constrained the speed of evolution of the observed characters drought adaptation and mycorrhizal strategy. Furthermore, it showed that this correlated evolution across the plant phylogeny was best characterized by a full or partial heterogeneous speed of evolution (Table 1). Finally, the calculated PIR scores for the different dataset versions were at least 20 times below the maximum recommended threshold of 0.1 (Table 3; Gardner & Organ, 2021), which ruled out potential issues of evolutionary sample size and phylogenetic imbalance, and provided strong support for the adequate detection of correlation evolution.

**Table 1.** Model rankings from the maximum-likelihood analysis of the evolutionary relationship between plant adaptation to drought and mycorrhizal strategy for each dataset version

| Evolution mode | Hidden rates | Model structure | *k* rates | Dataset v1 |  | Dataset v2 |  | Dataset v3 |  | Dataset v4 |  | Dataset v5 |  | Dataset v6 |  |
| --- | --- | --- | --- | --- | --- | --- | --- | --- | --- | --- | --- | --- | --- | --- | --- |
|  |  |  |  | AICc | AICcWt | AICc | AICcWt | AICc | AICcWt | AICc | AICcWt | AICc | AICcWt | AICc | AICcWt |
| Independent | 1 | ER | 2 | 2846.1 | < 0.01 | 2846.1 | < 0.01 | 2808.3 | < 0.01 | 2808.3 | < 0.01 | 1308.3 | < 0.01 | 1308.2 | < 0.01 |
| Independent | 1 | SYM | 4 | 2823.2 | < 0.01 | 2823.2 | < 0.01 | 2811.0 | < 0.01 | 2785.4 | < 0.01 | 1288.4 | < 0.01 | 1288.3 | < 0.01 |
| Independent | 1 | ARD | 8 | 2817.1 | < 0.01 | 2817.1 | < 0.01 | 2772.2 | < 0.01 | 2772.2 | < 0.01 | 1285.1 | < 0.01 | 1285.1 | < 0.01 |
| Independent | 2 | ER | 8 | 2684.4 | < 0.01 | 2684.3 | < 0.01 | 2619.5 | < 0.01 | 2619.6 | < 0.01 | 1234.6 | < 0.01 | 1234.6 | < 0.01 |
| Independent | 2 | SYM | 12 | 2651.7 | < 0.01 | 2651.7 | < 0.01 | 2586.9 | < 0.01 | 2586.9 | < 0.01 | 1217.9 | 0.01 | 1217.9 | 0.01 |
| Independent | 2 | ARD | 20 | 2625.9 | 0.04 | 2624.9 | 0.01 | 2589.3 | < 0.01 | 2568.3 | < 0.01 | 1215.1 | 0.03 | 1215.1 | 0.02 |
| Independent | 3 | ER | 18 | 2682.5 | < 0.01 | 2682.4 | < 0.01 | 2618.9 | < 0.01 | 2618.9 | < 0.01 | 1236.0 | < 0.01 | 1234.8 | < 0.01 |
| Independent | 3 | SYM | 24 | 2649.3 | < 0.01 | 2655.2 | < 0.01 | 2585.8 | < 0.01 | 2585.8 | < 0.01 | 1224.4 | < 0.01 | 1225.6 | < 0.01 |
| Independent | 3 | ARD | 36 | 2630.2 | < 0.01 | 2630.3 | < 0.01 | 2562.8 | < 0.01 | 2564.9 | < 0.01 | 1233.7 | < 0.01 | 1233.8 | < 0.01 |
| Dependent | 1 | ER | 1 | 4038.5 | < 0.01 | 4038.4 | < 0.01 | 3833.5 | < 0.01 | 3833.5 | < 0.01 | 1845.2 | < 0.01 | 1845.2 | < 0.01 |
| Dependent | 1 | SYM | 9 | 2764.0 | < 0.01 | 2764.0 | < 0.01 | 2753.0 | < 0.01 | 2753.0 | < 0.01 | 1258.8 | < 0.01 | 1258.8 | < 0.01 |
| Dependent | 1 | ARD | 18 | 2706.5 | < 0.01 | 2706.5 | < 0.01 | 2675.8 | < 0.01 | 2696.7 | < 0.01 | 1254.9 | < 0.01 | 1254.9 | < 0.01 |
| Dependent | 2 | ER | 4 | 3894.8 | < 0.01 | 3894.8 | < 0.01 | 3701.0 | < 0.01 | 3700.9 | < 0.01 | 1783.7 | < 0.01 | 1783.6 | < 0.01 |
| **Dependent** | **2** | **SYM** | **20** | 2659.4 | < 0.01 | 2659.4 | < 0.01 | 2596.0 | < 0.01 | 2596.0 | < 0.01 | **1207.8** | **0.97** | **1207.8** | **0.97** |
| **Dependent** | **2** | **ARD** | **38** | **2619.4** | **0.96** | **2616.3** | **0.99** | **2544.0** | **1.00** | **2553.6** | **1.00** | 1227.2 | < 0.01 | 1224.3 | < 0.01 |
| Dependent | 3 | ER | 9 | 3878.8 | < 0.01 | 3878.8 | < 0.01 | 3688.0 | < 0.01 | 3688.0 | < 0.01 | 1778.6 | < 0.01 | 1778.5 | < 0.01 |
| Dependent | 3 | SYM | 33 | 2636.7 | < 0.01 | 2643.8 | < 0.01 | 2588.2 | < 0.01 | 2592.6 | < 0.01 | 1222.1 | < 0.01 | 1222.2 | < 0.01 |
| Dependent | 3 | ARD | 60 | 2632.3 | < 0.01 | 2635.5 | < 0.01 | 2561.7 | < 0.01 | 2580.9 | < 0.01 | 1259.9 | < 0.01 | 1266.0 | < 0.01 |

ER, all rates are equal; SYM, only rates between any two states do not differ; ARD, all rates differ. The *k* rates is the number of evolutionary transition rate parameters independently estimated by a given model. The AICc is the sample size-corrected Akaike information criterion, while the AICcWt is the weighted AICc that provides the relative likelihood of each model per dataset version. The rows in bold are the best fitting model according to AICc for each dataset version. Dataset v1 and v2 were assembled with drought adaptation prioritizing DatasetID 92 of TRY and mycorrhizal strategy based on FungalRoot. Dataset v3 and v4 were assembled with drought adaptation prioritizing DatasetID 49 and 98 of TRY and mycorrhizal strategy based on FungalRoot. Dataset v5 and v6 were assembled with drought adaptation prioritizing DatasetID 92 of TRY and mycorrhizal strategy based on Bueno dataset. Odd and even dataset versions had their phylogenies mapped using the GBMB and GBOTB phylogeny, respectively. The assembly of dataset versions is summarized in Supporting Information Table S1 and described in the materials and methods.

**Table 2.** Ranking of the average rates for the evolutionary transitions from a mycorrhizal strategy and drought adaptation state to another.

| Hidden rate level | Mycorrhizal strategy | Drought adaptation |  | Mycorrhizal strategy | Drought adaptation | Transition rate (N°/100My) |
| --- | --- | --- | --- | --- | --- | --- |
| R1 | EEM | sensitive | → | EEM | tolerant | 26.636674973 |
| R1 | EEM | tolerant | → | EEM | sensitive | 22.593999567 |
| R2 | EEM | tolerant | → | EEM | sensitive | 11.494188976 |
| R2 | EEM | sensitive | → | EEM | tolerant | 3.582053911 |
| R1 | AM | sensitive | → | AM | tolerant | 2.650503866 |
| R1 | AM | tolerant | → | AM | sensitive | 1.634730831 |
| R1 | NR | tolerant | → | NR | sensitive | 0.177322608 |
| R1 | NR | sensitive | → | NR | tolerant | 0.033105750 |
| R2 | AM | tolerant | → | AM | sensitive | 0.008705943 |
| R2 | AM | sensitive | → | AM | tolerant | 0.006990306 |
| R2 | NR | sensitive | → | NR | tolerant | 0.004587933 |
| R2 | AM | sensitive | → | NR | sensitive | 0.002185387 |
| R2 | NR | tolerant | → | AM | tolerant | 0.002038248 |
| R2 | NR | sensitive | → | AM | sensitive | 0.001766620 |
| R2 | EEM | tolerant | → | NR | tolerant | 0.000594211 |
| R2 | AM | tolerant | → | NR | tolerant | 0.000425325 |
| R2 | AM | tolerant | → | EEM | tolerant | 0.000281040 |
| R1 | AM | sensitive | → | NR | sensitive | 0.000249479 |
| R2 | NR | tolerant | → | NR | sensitive | 0.000131191 |
| R1 | EEM | sensitive | → | AM | sensitive | 0.000123663 |
| R2 | NR | sensitive | → | EEM | sensitive | 0.000122614 |
| R1 | AM | tolerant | → | EEM | tolerant | 0.000121043 |
| R1 | NR | sensitive | → | AM | sensitive | 0.000115596 |
| R2 | EEM | tolerant | → | AM | tolerant | 0.000095959 |
| R1 | EEM | tolerant | → | AM | tolerant | 0.000087368 |
| R2 | NR | tolerant | → | EEM | tolerant | 0.000073183 |
| R1 | NR | sensitive | → | EEM | sensitive | 0.000065938 |
| R1 | AM | tolerant | → | NR | tolerant | 0.000050678 |
| R1 | AM | sensitive | → | EEM | sensitive | 0.000046256 |
| R2 | EEM | sensitive | → | AM | sensitive | 0.000036688 |
| R1 | EEM | sensitive | → | NR | sensitive | < 0.000000001 |
| R1 | EEM | tolerant | → | NR | tolerant | < 0.000000001 |
| R1 | NR | tolerant | → | AM | tolerant | < 0.000000001 |
| R1 | NR | tolerant | → | EEM | tolerant | < 0.000000001 |
| R2 | AM | sensitive | → | EEM | sensitive | < 0.000000001 |
| R2 | EEM | sensitive | → | NR | sensitive | < 0.000000001 |

**Table 3.**
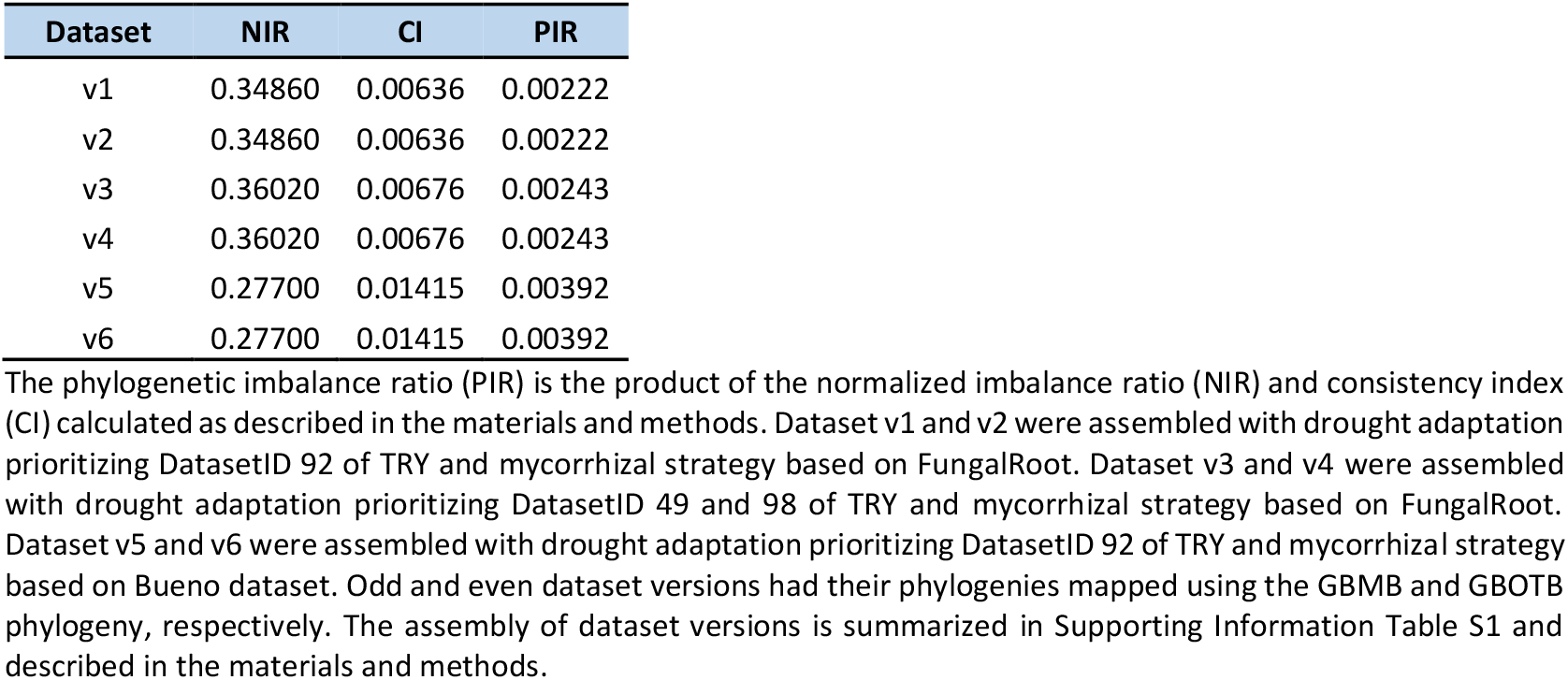
Phylogenetic imbalance ratio per dataset version.

### Mycorrhizas accelerate the evolution of plant adaptation to drought

My second aim was to characterize in detail the speed of evolution in terms of transition rates among the six composite states resulting from the combination of the two contrasting drought adaptations and the three different mycorrhizal strategies considered in this study. Hereto, I obtained the transition rate parameters estimated by the models best fitted to the six dataset versions, and analyzed their average values (Table 2; Supporting Information Table S3). In general, the top fastest transition rates were observed between the two contrasting plant adaptations to drought within a given mycorrhizal strategy, regardless of the hidden rate category (Table 2); the only exception to this was the rate of transitions from drought tolerance to sensitivity in lineages with the NR strategy in the hidden rate category two, which was slower than a few transitions among mycorrhizal strategies within a given drought adaptation (Table 2). Among the fastest transition rates, the top four occurred between drought sensitivity and tolerance and vice versa in lineages with the EEM strategy, regardless of the hidden rate category (Fig 2c; Table 2). These rates were on average – among all transition directions and hidden rate categories – approximately 15 and 300 times faster than that observed in lineages with the AM and NR strategy, respectively (Fig 2c; Table 2). This indicates that, at the evolutionary time scale, the speeds of these transitions were consistently superior in lineages that form ecto- or ericoid mycorrhizas, compared with that of the other mycorrhizal strategies tested here (Fig 2c; Table 2), confirming my second hypothesis. Moreover, the following fastest transition rates were observed between drought adaptations in the hidden rate category one in lineages with the AM strategy, which were higher than that in the lineages with the NR strategy in the same hidden rate category (Fig 2c; Table 2). These rates were tailed by a similar pattern in the hidden rate category two, with shifts between drought adaptations being faster in lineages with the AM strategy than in the lineages with the NR strategy (Fig 2c; Table 2). Taken together, this suggests that, although the hidden phylogenetic factor may influence the speed of gains and losses of drought tolerance, a consistent mycorrhizal association tend to accelerate the evolutionary processes of plant adaption to drought.

Another common aspect among lineages with either EEM or AM strategy was the impact of the hidden phylogenetic factor on the relationship between gains and losses of plant tolerance to drought. The hidden rate category one promoted more gains than losses of drought tolerance in lineages with either an AM or an EEM strategy, while the hidden rate category two had the opposite effect, i.e. it promoted more losses than gains of drought tolerance in lineages with either an AM or an EEM strategy (Table 2). In contrast, the lineages with the NR strategy showed an opposite pattern, with more losses than gains and more gains than losses of drought tolerance in the hidden rate category one and two, respectively (Table 2). By unveiling a consistent pattern of evolutionary influence common to lineages that persistently rely on mycorrhizal fungal symbionts, these results provide a closer understanding of the detected hidden phylogenetic factor. Furthermore, this factor also interacted markedly with the drought adaptation to influence the rate of evolutionary change in mycorrhizal strategy, with the top six fastest shifts occurring under the influence of the hidden rate category two (Table 2). Among these, the fastest rates were losses of AM strategy (i.e. transitions to NR strategy) in drought sensitive lineages, followed by gains of AM strategy (i.e. transitions from NR strategy) regardless of drought adaptation, which were then tailed by losses of EEM or AM strategy and symbiont shifts from AM to EEM strategy in drought tolerant lineages, respectively (Table 2). The relations among these transition rates were then altered by the hidden rate category one, with one main exception, i.e. like in the hidden rate category two, the fastest changes in mycorrhizal strategy in the hidden rate category one were also losses of AM strategy in drought sensitive linages (Table 2). Overall, these results help us to understand further the nature of the detected hidden phylogenetic factor, and unveiled that, independent of the latter, the most frequent and consistent changes in mycorrhizal strategy are AM losses in drought sensitive lineages.

### Sensitivity analysis showed consistency among main results

My main conclusions on the dependent evolution and on the differences among the mycorrhizal strategies in terms of transitions rates between drought adaptations were largely robust regarding the use of six partially different dataset versions (Table 1; Fig. 3). These dataset versions considered three sources of uncertainty, i.e. adaptation to drought data, mycorrhizal strategy data, and phylogenetic tree backbone (see materials and methods). Independently of the dataset version analyzed, the model ranking from the maximum-likelihood analysis consistently selected models of dependent evolution between the two plant characters, and always with two categories of hidden rates (Table 1). However, the structure of the best fitted model was slightly different among some of the dataset versions. While dataset v1 to v4 allowed for the best fitted model to have a structure with a fully heterogenous speed of evolution (ARD), the dataset v5 and v6 allowed for the best fitted model to have a structure with partially heterogenous speed of evolution (SYM) only (Table 1). This might be related to uncertainty in adaptation to drought and/or mycorrhizal strategy data, but it is not related to phylogenetic uncertainty. Moreover, because both dataset v5 and v6 had less than half of the sample size of the dataset v1 to v4 (Supporting Information Table S2), it is likely that dataset v5 and v6 limited in part the amplitude of the optimization process when attempting to estimate differences among all possible transition rates due to a restricted sample size.

**Fig. 3.**
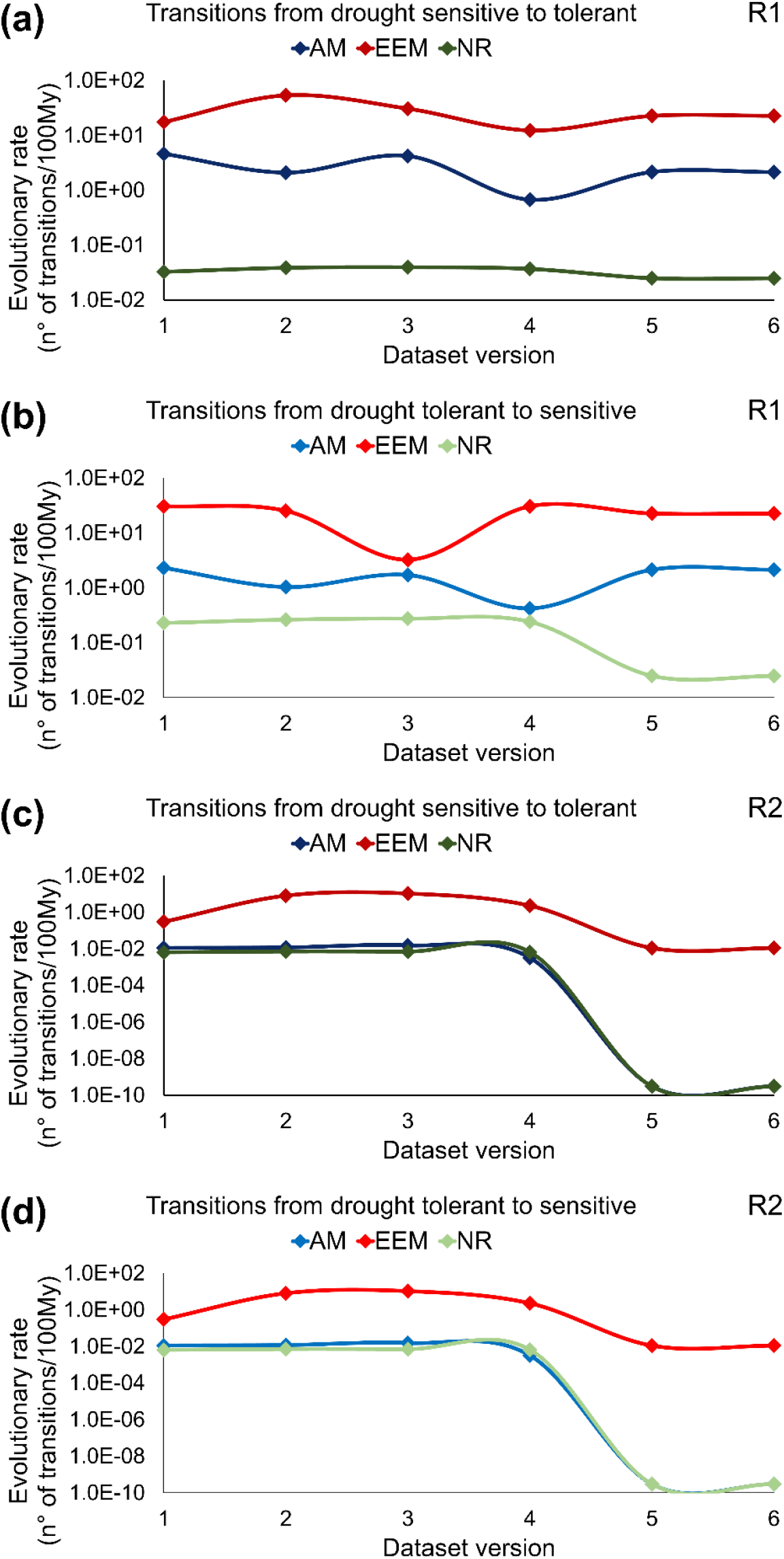
Sensitivity analysis on evolutionary transition rates between drought adaptations. Each panel shows the variation of evolutionary rates in logarithmic scale, as estimated by the models best fitted to dataset version 1 to 6, for the transitions from drought sensitive to tolerant (a) and tolerant to sensitive (b) in the hidden rate category one (R1), and from drought sensitive to tolerant (c) and tolerant to sensitive (d) in the hidden rate category two (R2), as influenced by the arbuscular mycorrhizal (AM), ecto- or ericoid mycorrhizal (EEM), or naked root (NR; non-mycorrhizal or facultatively AM) strategy. Dataset version 1 and 2 were assembled with drought adaptation prioritizing DatasetID 92 of TRY and mycorrhizal strategy based on FungalRoot. Dataset version 3 and 4 were assembled with drought adaptation prioritizing DatasetID 49 and 98 of TRY and mycorrhizal strategy based on FungalRoot. Dataset version 5 and 6 were assembled with drought adaptation prioritizing DatasetID 92 of TRY and mycorrhizal strategy based on Bueno dataset. Odd and even dataset versions had their phylogenies mapped using the GBMB and GBOTB phylogeny, respectively. The assembly of dataset versions is summarized in Supporting Information Table S1 and described in the materials and methods.

In spite of this minor structural variation, the main differences in transition rates between contrasting drought adaptations in lineages with different mycorrhizal strategies were largely consistent (Fig. 3). Regardless of the dataset version, the fastest transition rates were observed in the hidden rate category one (Fig. 3), and in this hidden rate category, the transition rates from drought sensitive to tolerant and vice versa had an increasing order of speed in lineages with NR, AM, and EEM strategy, respectively (Fig. 3a,b). A similar pattern was observed in the hidden rate category two when the dataset v1 to v3 were analyzed (Fig. 3b,c). Moreover, in the hidden rate category two, regardless of the dataset version analyzed the transitions rates from drought sensitive to tolerant and vice versa were also consistently faster in lineages with EEM strategy, compared with that of the other two mycorrhizal strategies (Fig. 3c,d). Only a minor deviance to the main pattern was observed for dataset v4 to v6 in the hidden rate category two (Fig. 3). In this hidden rate category, when dataset v4 was analyzed, the rate of transitions between drought adaptations was slightly slower in lineages with AM strategy, compared with that of the NR strategy, while when dataset v5 and v6 were analyzed, these rates were apparently similar (Fig. 3c,d). This provides a hint for a minor uncertainty in the impact of the hidden phylogenetic factor on the differences between AM and NR strategy in term of transition rates between drought adaptations. Among the changes in mycorrhizal strategy, the losses of AM strategy in drought sensitive lineages were consistent throughout dataset versions, except only in the hidden rate category one when dataset v4 was analyzed (Supporting Information Fig. S1). Moreover, the symbiont shifts from EEM to AM strategy in drought sensitive lineages were consistent throughout all dataset versions, except in the hidden rate category two when the dataset v3 was analyzed, while the other transitions were relatively more sensitive to data uncertainty depending on the hidden rate category (Supporting Information Fig. S1). Overall, the sensitivity analysis asserted the robustness of main conclusions on the dependent evolution between the observed plant characters, in which the evolutionary gains and losses of plant tolerance to drought consistently occurred at relatively faster rates in lineages that form the ecto- or ericod mycorrhizal strategy.

## Discussion

My results show that the evolutions of drought adaptation and mycorrhizal strategy in plants depend on each other, with the ecto- or ericoid mycorrhizal lineages as a group (EEM strategy) fast-tracking by 15 and 300 times more the evolutionary processes of plant adaption to drought, compared with that of the arbuscular mycorrhizal (AM) and naked root (NR, i.e. non-mycorrhizal or facultatively AM) strategy, respectively. This provides a quantitative demonstration that the evolution of a complex host phenotypic trait, which is essential to cope with critical environmental variation across temporal and spatial scales, is markedly shaped by host-microbe interactions.

Among ecological factors interacting with drought to influence vegetation shifts or stability, the most well studied are pests, pathogens, and grazing, while the role of beneficial biotic interactions are often neglected (reviewed by Martínez-Vilalta & Lloret, 2016), including mycorrhizas. My study, although based on an evolutionary timescale too coarse to be fully informative at finer ecological timescales, when combined with previous eco-physiological studies (Augé, 2001; Lehto & Zwiazek, 2011), supports the consideration of mycorrhizal strategy when studying the impact of droughts on vegetation to help us fully appreciate the mechanisms underpinning temporal variation and trajectory of responses. This would be particular important when considering vegetation types and larger time scales of post-drought monitoring, as it is increasingly recommended (Martínez-Vilalta & Lloret, 2016; Ploughe *et al*., 2019; Vilonen *et al*., 2022).

My results also raise the pressing question of what genetic, physiological, and ecological factors remain sufficiently stable at an evolutionary time scale to allow mycorrhizal plant linages to change more swiftly their adaptation to drought. These factors do not seem to be directly related with the abundance of genes of the symbiosis toolkit that many plant lineages carry across generations to form plant-microbe symbioses (Radhakrishnan *et al*., 2020). This is because ecto-, ericoid, and non-mycorrhizal plant lineages have higher degrees of symbiosis gene deletions compared with that of AM lineages (Radhakrishnan *et al*., 2020), while my results show that the rates of evolutionary transitions between drought adaptations are higher in the ecto-, ericoid, and AM lineages, compared to that of the non-mycorrhizal or facultatively AM lineages. Furthermore, symbiosis-related molecular pathways harbored in both mycorrhizal and non-mycorrhizal plants, such as the strigolactone and phenylpropanoid pathways (Fernández *et al*., 2019; Cosme *et al*., 2021; Venice *et al*., 2021), are unlikely to play a role on their own. However, single or combined symbiosis-related molecular pathways, whether mycorrhizal type-specific or not (Cope *et al*., 2019; Radhakrishnan *et al*., 2020), may contribute indirectly by allowing plants to control their mycorrhizal strategy, and in this way, provide access to physiological and ecological factors that may contribute more directly to shape the evolution of plant adaptation to drought.

Mycorrhizas are well known to influence the physiological capacity of plants to tolerate drought, with many studies documenting direct and indirect mechanisms, such as mycelial transport of water and nutrients, leading to beneficial effects on plant performance, including improved survival under drought (Augé, 2001; Lehto & Zwiazek, 2011; Sebastiana *et al*., 2018; Mu *et al*., 2021; Püschel *et al*., 2021). This would indicate that mycorrhizas might accelerate the evolutionary gains of drought tolerance by enhancing the transgenerational success of mycorrhizal plants that survive repeated droughts, compared with that of the less successful non-mycorrhizal counterparts. However, there are also many publications documenting no effects or negative effects of mycorrhizas on plant water relations during drought, particular for ecto-mycorrhizas, and it cannot be argued that mycorrhizas will always have a beneficial function (reviewed by Lehto & Zwiazek, 2011). In addition, my results suggest that the evolutionary advantage of mycorrhizas is more related to rapid adaptations rather than increases in tolerance only, by showing that mycorrhizas increase the gains of plant tolerance to drought nearly as much as the gains of sensitivity. These gains of sensitivity could be related, for instance, with the multifunctional roles of mycorrhizas combined with adaptation tradeoffs in plants (Van der Heijden *et al*., 2015; Pierce *et al*., 2017). According to the competitor, stress-tolerator, ruderal (CSR) theory, evolutionary gains of competitor and ruderal adaptations can only occur when lineages lose their stress-tolerance adaptation (Pierce *et al*., 2017). This mean that the ability to tolerate stress generally becomes a disadvantage in stably productive and frequently disturbed environments, where the other two contrasting adaptations tend to be more dominant, respectively (Li & Shipley, 2017; Pierce *et al*., 2017). Future studies combining my results with empirical testing could help us better understand how ecto-, ericoid, or arbuscular mycorrhizas influence the host survival and fitness in selective environments where drought sensitivity is a major eco-physiological advantage for the plants.

One plausible ecological explanation for how mycorrhizas facilitate the evolutionary processes of drought adaptation might be related to the imperfect vertical transmission of fidelity, recently modeled for host-microbiome associations (Bruijning *et al*., 2022). Although mycorrhizal fungi are not vertically transmitted, as are other microbes in other systems (Bruijning *et al*., 2022 and references therein), and plants have to reacquire fungi from soil every generation (Smith & Read, 2008), plant genomes harbor (or have lost) symbiosis-related genes that function as transgenerational transmission mechanisms of plant mycorrhizal strategy (Cope *et al*., 2019; Fernández *et al*., 2019; Radhakrishnan *et al*., 2020; Cosme *et al*., 2021; Venice *et al*., 2021). Moreover, it seems safe to assume that mycorrhizal fungal guilds are relatively stable entities at an evolutionary time scale based on fossil, chemistry, and genome observations (Remy *et al*., 1994; Redecker *et al*., 2000; Miyauchi *et al*., 2020; Huang *et al*., 2022). The transgenerational transmission of partner fidelity in the ecto- and ericoid mycorrhizal plant lineages seems to be relatively low, with about 6,000 ecto-mycorrhizal host species associating with more than 20,000 fungal species, and some of these fungi can form ericoid mycorrhizas with many plants (Walker *et al*., 2011; Van der Heijden *et al*., 2015). In contrast, the AM plant lineages seem to be relatively faithful, with more than 200,000 host species associating with about 300 fungal species (Van der Heijden *et al*., 2015; Davison *et al*., 2020). Non-mycorrhizal or facultatively AM plants either do not associate or associate to a much lesser extent with mycorrhizal fungi (Brundrett & Tedersoo, 2018a; Cosme *et al*., 2018). Functional variation of mycorrhizal effects on plant water relations has been documented (Augé, 2001; Lehto & Zwiazek, 2011), and is particularly obvious for ecto-mycorrhizas, where mycelium exploration types vary from close contact explorations, with smooth mycorrhizal tips having only a few short emanating hyphae, to extremely long explorative distances with highly differentiated rhizomorphs (Lehto & Zwiazek, 2011; Ekblad *et al*., 2013). Hence, a low host fidelity across plant generations might enhance offspring phenotypic variance due to the functional variation among mycorrhizal fungi. Under a fluctuating natural selection, due to changes in water availability over time, this enhanced offspring phenotypic variance is likely to ensure that at least some plant individuals are able to maintain a non-zero fitness in any given time step, lowering the likelihood of extinction, and increasing the rate of adaptation (see general model by Bruijning *et al*., 2022). This could potentially explain how the contrasting mycorrhizal strategies differently shape the speed of evolution of drought adaptation in plants.

In sum, my study indicates that the evolution of plant adaptation to critical environmental variations in water availability is inherently dependent on mycorrhizas. This embodies a previously overlooked, yet important long-term advantage of host-microbe symbiosis with ecological consequences of global proportions, and could be linked to imperfect transgenerational transmission of partner fidelity. Future investigations of this and related working hypotheses under past and future climate change scenarios are absolutely indispensable to yield important insights into the role of mycorrhizas in plant evolution.

## Acknowledgements

MC was supported by the European Commission’s grant H2020-MSCA-IF-2018 ‘SYMBIO-INC’ (GA 838525)”. Computational resources were provided by the supercomputing facilities of the Université catholique de Louvain (CISM/UCL) and the Consortium des Équipements de Calcul Intensif en Fédération Wallonie Bruxelles (CÉCI) funded by the Fond de la Recherche Scientifique de Belgique (F.R.S.-FNRS) under convention 2.5020.11 and by the Walloon Region. The author is thankful to Stéphane Declerck for office space and to Damien François, James Boyko, and Jacob Gardner for methodological clarifications.

## Author Contributions

MC designed the research, collected and analyzed the data, interpreted the results, and wrote the manuscript.

## Data Availability

The main Figures and Tables are available in the article. The Supporting Information Figures and Tables are available in the supporting information of the article. The Supporting Information Data and Note are available in a publicly repository at https://github.com/marcosmeweb/symbio-inc-drought.

## Supporting Information

**Fig. S1** Sensitivity analysis on the rates of evolutionary transitions among mycorrhizal strategies.

**Table S1** Summary of the assembly of the six dataset versions analyzed.

**Table S2** Number of plant species grouped by drought adaptation and mycorrhizal strategy according to each dataset version.

**Table S3** Evolutionary transition rates estimated by the best fitted model for each dataset version.

**Data S1** Files of phylogenetic trees and data frames with plant mycorrhizal strategy and drought adaptation assignments according each dataset version.

**Data S2** File with the geographical occurrences for 1,066 of the species included in the analysis.

**Data S3** Output files of all corHMM runs.

**Note S1** R script with the codes used to run all hidden Markov models.

## Notes

https://github.com/marcosmeweb/symbio-inc-drought

### Competing Interest Statement

The authors have declared no competing interest.

