## Supporting Information for "Mycorrhizas shape the evolution of plant adaptation to drought"

Article title: Mycorrhizas shape the evolution of plant adaptation to drought
Authors: Marco Cosme
Article acceptance date: NA

The following Supporting Information is available for this article:

**Fig. S1** Sensitivity analysis on the rates of evolutionary transitions among mycorrhizal strategies.

**Table S1** Summary of the assembly of the six dataset versions analyzed.

**Table S2** Number of plant species grouped by drought adaptation and mycorrhizal strategy according to each dataset version.

**Table S3** Evolutionary transition rates estimated by the model best fitted to each dataset version.

**Data S1** Files of phylogenetic trees and data frames with plant mycorrhizal strategy and drought adaptation assignments according each dataset version.

**Data S2** File with the geographical occurrences for 1,066 of the species included in the analysis.

**Data S3** Output files of all corHMM runs.

**Notes S1** R script with the codes used to run all hidden Markov models.

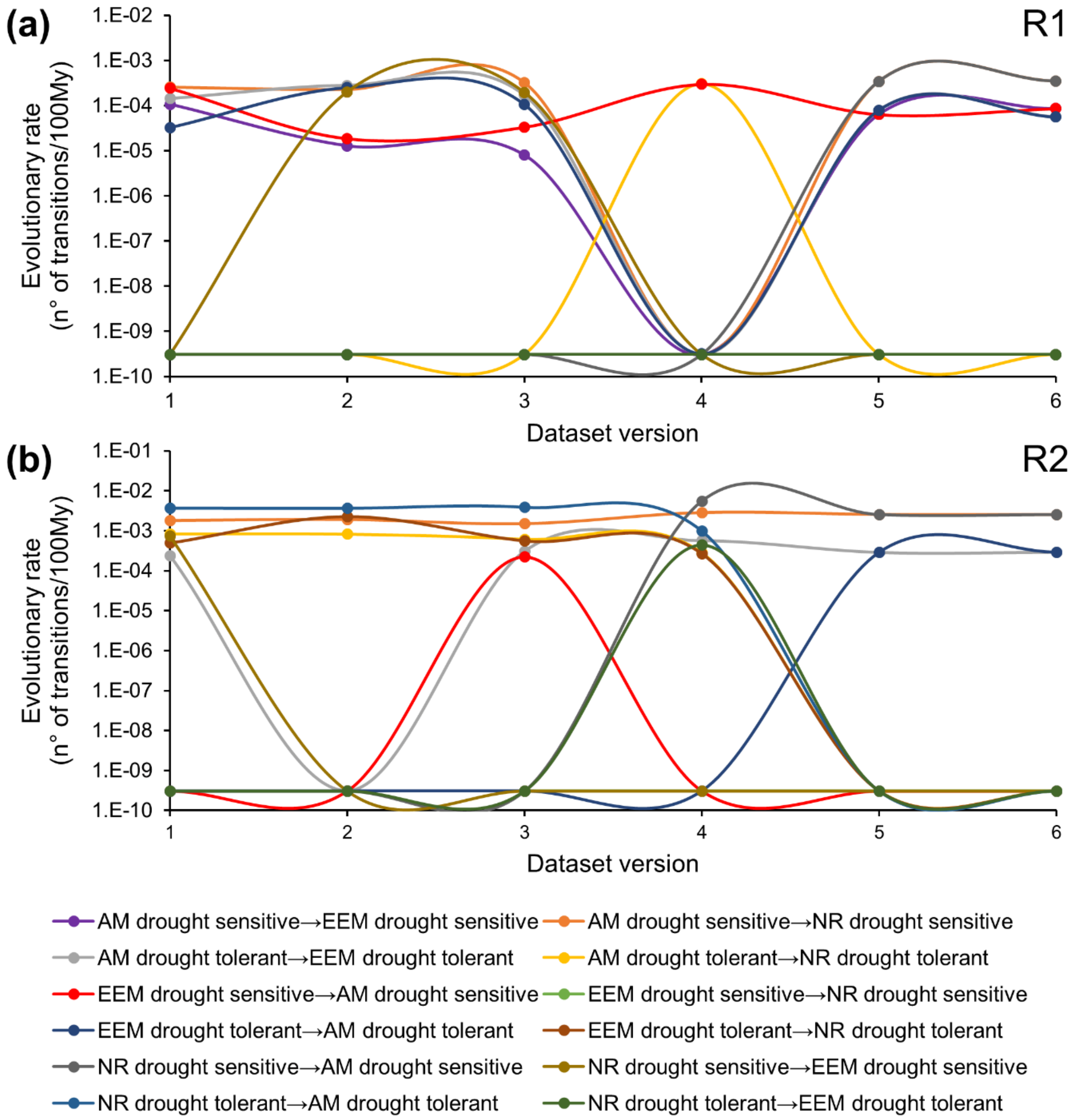


**Fig. S1** Sensitivity analysis on the rates of evolutionary transitions among mycorrhizal strategies. (a) and (b) show the evolutionary transition rates from the arbuscular mycorrhizal (AM), ecto- or ericoid mycorrhizal (EEM), or naked root (NR; non-mycorrhizal or facultatively AM) strategy to a different mycorrhizal strategy in lineages that have either a drought sensitive or tolerant adaptation in the hidden rate category one (R1) and two (R2), respectively. These rates were estimated by the best fitted model for the dataset version 1 to 6. Differences among dataset versions are summarized in Supporting Information Table S1 and described in the materials and methods.

**Table S1** Summary of the assembly of the six dataset versions analyzed.

| Dataset | Drought adaptation | Mycorrhizal strategy | Phylogenetic tree |
|---|---|---|---|
| **v1** | Prioritizing DatasetID 92 | Based on FungalRoot | Phylogeny mapped against GBMB |
| **v2** | Prioritizing DatasetID 92 | Based on FungalRoot | Phylogeny mapped against GBOTB |
| **v3** | Prioritizing DatasetID 49 and 98 | Based on FungalRoot | Phylogeny mapped against GBMB |
| **v4** | Prioritizing DatasetID 49 and 98 | Based on FungalRoot | Phylogeny mapped against GBOTB |
| **v5** | Prioritizing DatasetID 92 | Based on Bueno dataset | Phylogeny mapped against GBMB |
| **v6** | Prioritizing DatasetID 92 | Based on Bueno dataset | Phylogeny mapped against GBOTB |

DatasetID refers to the number of the original dataset made available by TRY - plant trait database (Kattge et al., 2020). FungalRoot refers to the dataset published by Soudzilovskaia et al. (2020) while Bueno dataset refers to the dataset published by Bueno et al. (2018). GBMB and GBOTB refers to the plant phylogeny versions published by Smith & Brown (2018).

**Table S2** Number of plant species grouped by drought adaptation and mycorrhizal strategy according to each dataset version.

| Drought adaptation | Mycorrhizal strategy | Dataset | | |
|---|---|---|---|---|
| | | v1 or v2 | v3 or v4 | v5 or v6 |
| Sensitive | AM | 408 | 390 | 232 |
| | EEM | 169 | 157 | 83 |
| | NR | 181 | 182 | 110 |
| Tolerant | AM | 637 | 655 | 238 |
| | EEM | 177 | 189 | 104 |
| | NR | 66 | 65 | 20 |
| Total | | 1638 | 1638 | 787 |

AM, arbuscular mycorrhizal; EEM, ecto- or ericoid mycorrhizal; NR, naked root, i.e. non-mycorrhizal or facultatively AM. The assembly of dataset versions is summarized in Supporting Information Table S1 and described in the materials and methods.

**Table S3** Evolutionary transition rates estimated by the model best fitted to each dataset version.

| Hidden rate level | Mycorrhizal strategy | Drought adaptation | | Mycorrhizal strategy | Drought adaptation | Evolutionary rate (n° transitions/100My) | | | | | | |
|---|---|---|---|---|---|---|---|---|---|---|---|---|
| | | | | | | Dataset v1 | Dataset v2 | Dataset v3 | Dataset v4 | Dataset v5 | Dataset v6 | Average |
| R1 | AM | sensitive | → | AM | tolerant | 4.6178226164 | 2.0992073495 | 4.2131428417 | 0.6752361653 | 2.1534487333 | 2.1441654881 | 2.6505038657 |
| R1 | AM | sensitive | → | EEM | sensitive | 0.0001073912 | 0.0000128746 | 0.0000080641 | 0.0000000003 | 0.0000633072 | 0.0000858977 | 0.0000462558 |
| R1 | AM | sensitive | → | NR | sensitive | 0.0002546187 | 0.0002271891 | 0.0003214892 | 0.0000000003 | 0.0003427322 | 0.0003508438 | 0.0002494789 |
| R1 | AM | tolerant | → | AM | sensitive | 2.3220188933 | 1.0386175227 | 1.7261406781 | 0.4239936719 | 2.1534487333 | 2.1441654881 | 1.6347308312 |
| R1 | AM | tolerant | → | EEM | tolerant | 0.0001420844 | 0.0002793294 | 0.0001698615 | 0.0000000003 | 0.0000791728 | 0.0000558117 | 0.0001210433 |
| R1 | AM | tolerant | → | NR | tolerant | 0.0000000003 | 0.0000000003 | 0.0000000003 | 0.0003040686 | 0.0000000003 | 0.0000000003 | 0.0000506784 |
| R1 | EEM | sensitive | → | AM | sensitive | 0.0002440739 | 0.0000184770 | 0.0000331782 | 0.0002970445 | 0.0000633072 | 0.0000858977 | 0.0001236631 |
| R1 | EEM | sensitive | → | EEM | tolerant | 17.5624335452 | 53.7188431943 | 30.7644977696 | 12.3354825312 | 22.7142312774 | 22.7245615189 | 26.6366749728 |
| R1 | EEM | sensitive | → | NR | sensitive | 0.0000000003 | 0.0000000003 | 0.0000000003 | 0.0000000003 | 0.0000000003 | 0.0000000003 | 0.0000000003 |
| R1 | EEM | tolerant | → | AM | tolerant | 0.0000327271 | 0.0002499864 | 0.0001065126 | 0.0000000003 | 0.0000791728 | 0.0000558117 | 0.0000873685 |
| R1 | EEM | tolerant | → | EEM | sensitive | 30.7644977696 | 25.3393806624 | 3.2568284023 | 30.7644977696 | 22.7142312774 | 22.7245615189 | 22.5939995667 |
| R1 | EEM | tolerant | → | NR | tolerant | 0.0000000003 | 0.0000000003 | 0.0000000003 | 0.0000000003 | 0.0000000003 | 0.0000000003 | 0.0000000003 |
| R1 | NR | sensitive | → | AM | sensitive | 0.0000000003 | 0.0000000003 | 0.0000000003 | 0.0000000003 | 0.0003427322 | 0.0003508438 | 0.0001155962 |
| R1 | NR | sensitive | → | EEM | sensitive | 0.0000000003 | 0.0002005087 | 0.0001951204 | 0.0000000003 | 0.0000000003 | 0.0000000003 | 0.0000659384 |
| R1 | NR | sensitive | → | NR | tolerant | 0.0328052837 | 0.0388376070 | 0.0398098750 | 0.0371048582 | 0.0251032600 | 0.0249736145 | 0.0331057497 |
| R1 | NR | tolerant | → | AM | tolerant | 0.0000000003 | 0.0000000003 | 0.0000000003 | 0.0000000003 | 0.0000000003 | 0.0000000003 | 0.0000000003 |
| R1 | NR | tolerant | → | EEM | tolerant | 0.0000000003 | 0.0000000003 | 0.0000000003 | 0.0000000003 | 0.0000000003 | 0.0000000003 | 0.0000000003 |
| R1 | NR | tolerant | → | NR | sensitive | 0.2306874607 | 0.2646362426 | 0.2762978020 | 0.2422372695 | 0.0251032600 | 0.0249736145 | 0.1773226082 |
| R2 | AM | sensitive | → | AM | tolerant | 0.0112881356 | 0.0119393993 | 0.0154691346 | 0.0032451682 | 0.0000000003 | 0.0000000003 | 0.0069903064 |
| R2 | AM | sensitive | → | EEM | sensitive | 0.0000000003 | 0.0000000003 | 0.0000000003 | 0.0000000003 | 0.0000000003 | 0.0000000003 | 0.0000000003 |
| R2 | AM | sensitive | → | NR | sensitive | 0.0017992134 | 0.0019057686 | 0.0014967186 | 0.0028298916 | 0.0025432399 | 0.0025374896 | 0.0021853869 |
| R2 | AM | tolerant | → | AM | sensitive | 0.0150617890 | 0.0170584682 | 0.0201154011 | 0.0000000003 | 0.0000000003 | 0.0000000003 | 0.0087059432 |
| R2 | AM | tolerant | → | EEM | tolerant | 0.0002376114 | 0.0000000003 | 0.0003085181 | 0.0005643560 | 0.0002863425 | 0.0002894119 | 0.0002810400 |
| R2 | AM | tolerant | → | NR | tolerant | 0.0008352891 | 0.0008236262 | 0.0006128015 | 0.0002802334 | 0.0000000003 | 0.0000000003 | 0.0004253251 |
| R2 | EEM | sensitive | → | AM | sensitive | 0.0000000003 | 0.0000000003 | 0.0002201288 | 0.0000000003 | 0.0000000003 | 0.0000000003 | 0.0000366884 |
| R2 | EEM | sensitive | → | EEM | tolerant | 0.3061230154 | 8.1793437785 | 10.6737577044 | 2.3105497272 | 0.0112528002 | 0.0112964414 | 3.5820539112 |
| R2 | EEM | sensitive | → | NR | sensitive | 0.0000000003 | 0.0000000003 | 0.0000000003 | 0.0000000003 | 0.0000000003 | 0.0000000003 | 0.0000000003 |
| R2 | EEM | tolerant | → | AM | tolerant | 0.0000000003 | 0.0000000003 | 0.0000000003 | 0.0000000003 | 0.0002863425 | 0.0002894119 | 0.0000959593 |
| R2 | EEM | tolerant | → | EEM | sensitive | 0.0484047170 | 38.0082268810 | 30.7644977696 | 0.1214552458 | 0.0112528002 | 0.0112964414 | 11.4941889758 |
| R2 | EEM | tolerant | → | NR | tolerant | 0.0004909987 | 0.0022530685 | 0.0005543760 | 0.0002668236 | 0.0000000003 | 0.0000000003 | 0.0005942112 |
| R2 | NR | sensitive | → | AM | sensitive | 0.0000000003 | 0.0000000003 | 0.0000000003 | 0.0055189926 | 0.0025432399 | 0.0025374896 | 0.0017666205 |
| R2 | NR | sensitive | → | EEM | sensitive | 0.0007356844 | 0.0000000003 | 0.0000000003 | 0.0000000003 | 0.0000000003 | 0.0000000003 | 0.0001226143 |
| R2 | NR | sensitive | → | NR | tolerant | 0.0065125802 | 0.0072559823 | 0.0071219358 | 0.0066371009 | 0.0000000003 | 0.0000000003 | 0.0045879333 |
| R2 | NR | tolerant | → | AM | tolerant | 0.0036969122 | 0.0036675912 | 0.0038755436 | 0.0009894432 | 0.0000000003 | 0.0000000003 | 0.0020382485 |
| R2 | NR | tolerant | → | EEM | tolerant | 0.0000000003 | 0.0000000003 | 0.0000000003 | 0.0004390984 | 0.0000000003 | 0.0000000003 | 0.0000731833 |
| R2 | NR | tolerant | → | NR | sensitive | 0.0000000003 | 0.0000000003 | 0.0005352982 | 0.0002518464 | 0.0000000003 | 0.0000000003 | 0.0001311910 |

Hidden rate level corresponds to the level (R1 or R2) of the hidden phylogenetic factor that influences a given transition. AM, arbuscular mycorrhizal; EEM, ecto- or ericoid mycorrhizal; NR, naked root, i.e. non-mycorrhizal or facultatively AM. A summary of the dataset versions is provided in the Supporting Information Table S1 and the assembly of dataset versions is described in materials and methods. Rows colored in grey correspond to transitions between drought adaptations within a given mycorrhizal strategy. Rows colored in white correspond to transitions among mycorrhizal strategies within a given drought adaptation.

**Data S1** Files of phylogenetic trees and data frames with plant mycorrhizal strategy and drought adaptation assignments according each dataset version.
Found at https://github.com/marcosmeweb/symbio-inc-drought

**Data S2** File with the geographical occurrences for 1,066 of the species included in the analysis.
Found at https://github.com/marcosmeweb/symbio-inc-drought

**Data S3** Output files of all corHMM runs.
Found at https://github.com/marcosmeweb/symbio-inc-drought

**Note S1** R script with the codes used to run all hidden Markov models.
Found at https://github.com/marcosmeweb/symbio-inc-drought
